# A multilamellar organelle for chemosymbiosis in an aplacophoran mollusc adapted to anoxic cold seep sediment

**DOI:** 10.1101/2025.01.16.633305

**Authors:** Yunlong Li, Chong Chen, Xu Liu, Menggong Li, Haiyun Zhou, Hui Wang, Xiaoshan Zheng, Xing He, Shanshan Liu, Jian-Wen Qiu, Pei-Yuan Qian, Jin Sun

## Abstract

Symbiosis with chemoautotrophic bacteria has evolved in many animal lineages inhabiting reducing habitats such as hydrothermal vents, allowing these holobionts to thrive in dark biospheres^1^. In certain instances, the symbionts have become intracellular, residing within specialised bacteriocytes^2^. The integration of microbial symbionts with eukaryotic cells vary across known animals; however, no specialised organelle dedicated to chemosynthesis has been identified yet^2^. Here, we report a mode of symbiosis where sulphur-oxidising bacteria cultured within spherical multilamellar compartments (~12 μm) in the cold-seep aplacophoran mollusc *Chaetoderma shenloong*. This organelle, which we name ‘dracosphera’, is ubiquitous within the hypertrophied and intricately reticulate digestive gland of *C. shenloong*, which has otherwise lost most of its gut. Given that the symbionts are strictly anaerobic and the host resides in anoxic sediments tens of centimetres below the surface, the dracosphera may serve to minimise oxygen diffusion to the bacteria, akin to mechanisms observed in microbial diazotrophs^3^ or termite hindguts^4^, as supported by our genomic and spatial transcriptomic analyses. Hypoxic conditions have been known to induce radical adaptations in meiofauna, exemplified by the acquisition of hydrogenosomes^5^. Our discovery of similarly exceptional adaptations in *C. shenloong* provides new insights into the evolution of such organelles also in larger animals.

## Main Text

Symbiotic interactions are prevalent across the Tree of Life, occurring at various levels of intimacy^6^. The success of eukaryotic life is largely attributed to the energy conferred by symbiosis, which ultimately led to the acquisition of organelles such as mitochondria and chloroplasts^7,8^. A major scientific finding following the discovery of the first deep-sea hydrothermal vent on the Galápagos Rift in 1977^9^ was that many animals inhabiting reducing habitats, such as hot vents and cold seeps, derive their energy and nutrients by ‘feeding’ on symbiotic bacteria that perform chemosynthesis utilising inorganic chemicals^10^. Today, we recognise chemosymbiosis as widespread throughout the ocean and playing a pivotal role in forming the highly productive vent and seep ecosystems^1,2^, with new and unexpected holobionts continuing to be discovered^11,12^.

Mode of chemosymbiosis in large-bodied animals varies significantly. Some species cultivate and derive nutrition from epibiotic bacteria, as observed in various crustaceans^13,14^. Others occupy a transitional zone between extracellular and intracellular, where the symbionts reside within host vacuoles but maintain connections to the external environment, such as bathymodioline mussels and abyssochrysoid snails^15,16^. Additionally, some lineages exhibit complete encapsulation of symbionts by host bacteriocyte membranes, as seen in siboglinid tubeworms and peltospirid snails^10,12,17^. While the structures of symbiotic organs differ, most lack further compartmentalisation of symbionts within the bacteriocytes^12,18,19^, except for lucinid bivalves which house them inside vacuoles^20^. To date, no organelle dedicated to chemosynthesis has been identified.

Recently, a giant aplacophoran worm-mollusc from the understudied class Caudofoveata^21^ was collected from black, reducing sediments of the Haima deep-sea cold seep in the South China Sea. This species, named *Chaetoderma shenloong* (‘divine dragon’ in Chinese)^22,23^, exhibits an exceptional size, reaching over 150 mm, in contrast to most species in the class, which average around 10 mm^24^. Despite its remarkable size and unique habitat—buried approximately 40 cm below the sediment surface in black, reducing mud adjacent to a colony of symbiotic vesicomyid clams (Fig. 1a; Extended Data Fig. 1) — little is known about its ecology. The presence of multiple individuals within a single 6.5 cm diameter push-core in diameter (Extended Data Fig. 1) indicates that *C. shenloong* forms dense aggregations within the sediment, which becomes completely anoxic just 3 mm below the surface (Extended Data Fig. 1). This distribution is markedly different from typical caudofoveates, which usually burrow only in the upper 2-3 cm of the sediment^25^. Such high densities of large-bodied animals at chemosynthetic habitats is a hallmark of symbiotic species^26^. This raises the question: could *C. shenloong* be a holobiont, and if so, where do the symbionts reside? Here, we address this question using a variety of techniques, including traditional dissection, three-dimensional anatomical reconstruction with μ-computed tomography (CT), fluorescent *in situ* hybridisation (FISH), electron microscopy, holobiont genomics, and spatial transcriptomics, revealing *C. shenloong* to be the first animal with an organelle dedicated to chemosymbiosis.

**Fig. 1.**
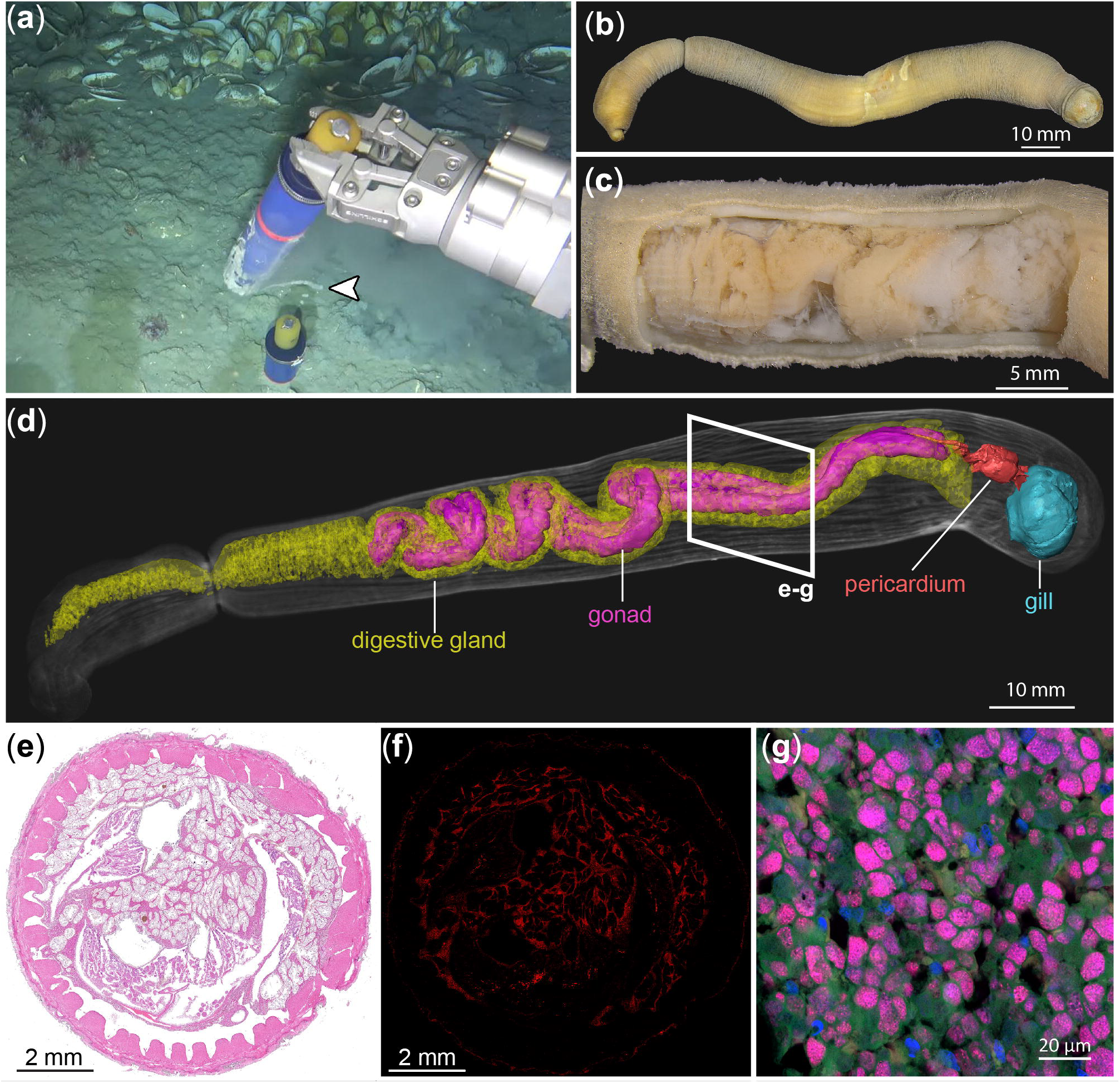
**a**, *In situ* imagery of *Chaetoderma shenloong* while being sampled from deep sediments around a colony of vesicomyid clams using a push-corer. **b**, External morphology of *C. shenloong*. **c**, The trunk with the body wall dissected away to show the symbiotic organ (digestive gland). **d**, 3D reconstruction of the key internal organs from μ-CT scan. **e**, Histological section stained with hematoxylin and eosin. **f**, Spatial distribution of symbionts in the section shown by fluorescence in situ hybridization (FISH) of a symbiont-specific probe (red signal). **g**, A close-up FISH imagery localising symbionts within spherical structures (‘dracosphera’); showing symbiont (red), nucleus (blue), membrane (green), and eubacteria (cyan).

### Anatomical and Stable Isotopic Evidence

Although the external morphology of *C. shenloong* (Fig. 1b) does not diverge greatly from typical caudofoveates except for its large size and broad body^22^, dissection of *C. shenloong* (Fig. 1c) revealed peculiar internal anatomy where most of the body cavity from the anterium throughout the trunk (demarcated by a constricted ‘neck’) is taken up by what seems to be a single coiled organ with haemocoel (blood) filling up the rest. Both 3D reconstruction from μ-CT scanning (Fig. 1d) and sectioning (Fig. 1e) showed this to be a complex, reticulated mesh-like organ – interpreted to be a greatly hypertrophied and modified digestive gland (diverticulum) due to its anterior connection with the buccal mass containing a very reduced radula^22^ – wrapped around a well-developed gonad. Much of the digestive tract from the oesophagus to stomach to the intestine appears to have been lost in *C. shenloong*, which is present in typical omnivorous caudofoveates feeding on detritus, foraminifera, or small animals^25^. Reduction or loss of the digestive tract is typical among chemosymbiotic holobionts^27^. The posterium is more typical of the class, containing a pericardium and a sizeable gill opening to the mantle cavity.

Our FISH results (Fig. 1e-g and Extended Data Fig. 2) combining general and symbiont-specific bacterial probes revealed strongly localised chemosymbiotic bacterial signals in the digestive gland of *C. shenloong*, providing the first piece of evidence that this organ is symbiotic. Using the digestive gland as a symbiotic organ has independently evolved in several molluscan lineages, most notably the extensive zooxanthellate tubular system in *Tridacna* giant clams for housing algal photosymbionts^28^ and the ‘solar-powered’ slug *Elysia* which uses digestive tubules to contain chloroplasts of a kleptoplastic origin^29,30^. This is also analogous to the symbiotic organs of the vent peltospirid snails *Chrysomallon* and *Gigantopelta*, which are modified oesophageal glands and therefore also originate from the digestive system^12,17^. Stable isotope ratios of carbon (δ^13^C) and nitrogen (δ^15^N) also lend support to *C. shenloong* relying on sulphur-oxidising symbiosis (Extended Data Fig. 3a), when compared to other fauna from the same seep^31^. A close examination of the digestive gland of *C. shenloong* using confocal microscopy (Fig. 1f) showed that the symbiont signals are restricted to small, spherical structures between 10-15 μm in diameter densely populating the reticulated diverticulum (Extended Data Fig. 2j-k).

From scanning electron microscopy (SEM) of the digestive gland (Fig. 2a), we consider these spherical structures to be intracellular as they are wrapped inside a layer of membranous structure, most likely the cell membrane of the host’s bacteriocyte. However, the complex nature of membranous structures within the symbiotic organ under transmission electron microscopy (TEM) makes the interpretation of cell boundaries challenging (Extended Data Fig. 4). These spherical structures are multilamellar and densely packed in bacterial cells under TEM (Fig. 2b), a mode of symbiont structural integration unlike any other known chemosymbiosis^2^. We here name this previously undocumented multilamellar organelle as the ‘dracosphera’ – a combination of *draco* meaning ‘dragon’ and *sphera* meaning ‘ball, sphere’ in Latin. The dracospherae exhibited more than 20 layers of membranes (100-1000 nm thick). Our serial imaging of a complete dracosphera using focused-ion beam scanning electron microscopy (FIB-SEM) combined with FISH confirmed the absence of eukaryotic nucleus and lysosomes within the dracosphera but small vacuoles with a single layer of membrane were present (Extended Data Fig. 4). Though smaller multilamellar bodies have been seen in the giant tubeworm *Riftia*, those were much smaller and are linked to symbiont digestion^19^.

**Fig. 2.**
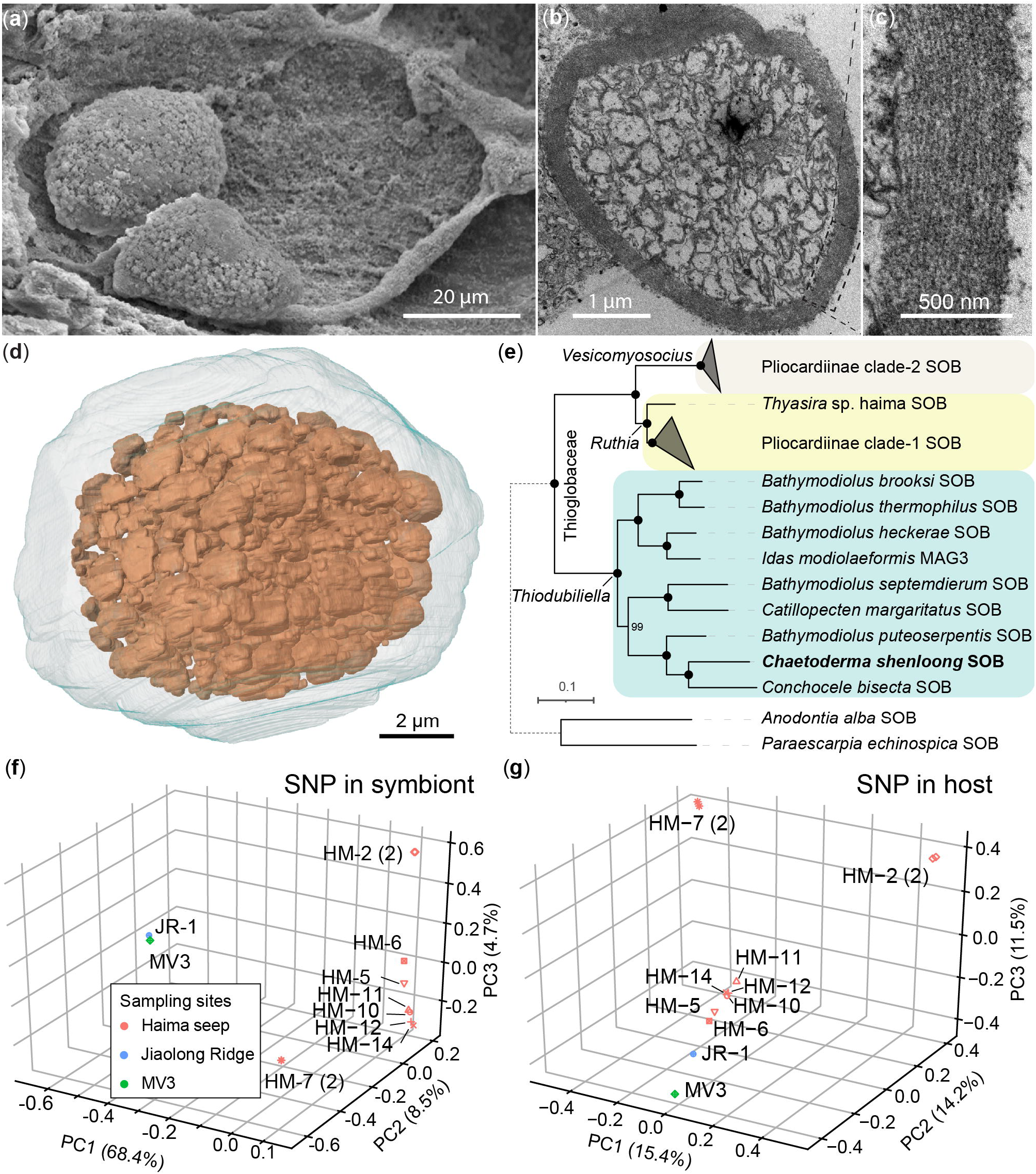
**a**, SEM micrograph of two dracospherae enveloped by a membrane. **b**, TEM micrograph showing cross-section view of a dracosphera. **c**, Close-up of the mutilamellar wall of the same dracosphera. **d**, 3D reconstruction of a dracosphera from FIB-SEM data showing tightly-packed symbiont cells completely enveloped. **e**, Phylogenetic position of the *Chaetoderma shenloong* symbiont (IQ-TREE 2, best-fitting model with partition). **f**, Principal component analysis (PCA) of symbiont SNPs plotted using the top three components. **g**, The same for host SNPs.

### Molecular Characterisation of the Symbiosis

A single sulphur-oxidising bacteria (SOB) symbiont species in the genus *Thiodubiliella* was found inside the dracosphera, up to 1629 symbiont cells per eukaryotic cell within the reticulated mesh-like digestive diverticula (Extended Fig. 5d). It encodes only 1022 genes in a 1.14 Mb genome that is more reduced than its close relatives (*Bathymodiolus* and *Conchocele bisecta* SOB symbionts, Extended Data Fig. 5a-b). Three genes are absent in the TCA cycle and two in the biosynthesis of amino acids (Methionine and Tyrosine), which are compensated by the host (Extended Data Fig. 6). Remarkably, the genomic and transcriptomic analyses showed that the symbiont is capable of facultative anaerobic life, actively expressing genes in dissimilatory nitrate reduction that could play vital roles as the electron acceptor (Extended Data Fig. 6). The loss of the *caa*_*3*_*-*type cytochrome oxidase^32^, combined with the presence of the *cbb*_3_-type with a high affinity for oxygen under hypoxia, indicates this SOB is adapted to hypoxic environments^33-35^.

We examined the genetic divergence patterns of both the host and symbiont using single nucleotide polymorphisms (SNPs), where a congruent pattern suggests vertical transmission while a conflicting pattern would indicate horizontal transmission. The identical symbiont SNPs observed in the anterior and posterior digestive gland of the same host individual (Fig. 2g and Extended Data Fig. 5c) suggests a limited time window for the symbiont acquisition. Among SNPs from eight hosts collected from Haima (Fig. 2g), two individuals (‘HM-2’, ‘HM-7’) were divergent from the rest which also had symbiont SNPs that differed from the others. We also detected some symbiont signals in the gonad through FISH (Fig. 1f). Overall, these lend support to a vertical transmission of the symbionts, which is consistent with the very small genome size of the symbiont. Nevertheless, a mixed mode of symbiont acquisition like solemyid clams^36^ and *Chrysomallon* snails^37^ cannot be ruled out.

### Host and symbiont interactions

As a reference, we assembled a new chromosome-level genome of *C. shenloong* from Haima, updating from an existing one collected from Jiaolong Ridge, also in the South China Sea^23^. The co-localisation of symbiotic reads and an unsupervised cluster of host reads highlights the potential to investigate the host and symbiont interaction, indicated by the spatial (meta)transcriptomics (Fig. 3a-c and Extend Data Fig. 7). Under the aggregated bin 20 (20 × 20 DNBs, i.e. 10 μm diameter), the average numbers of the captured molecular identifiers (MID) and genes per bin in 5 sections were 56 and 23, respectively (Supplementary file Table S7). There was a significantly higher ratio of symbiont-derived reads in symbiotic clusters compared to the rest, up to 45% vs. 27% in chip-5 (*p* < 0.0001, Fig. 3d). In total, 253 candidate genes were selected with the threshold as marker genes across at least two independent sections (Fig. 3e, labelled in bold) and then further classified on their functions (Fig. 3f-g).

**Fig. 3.**
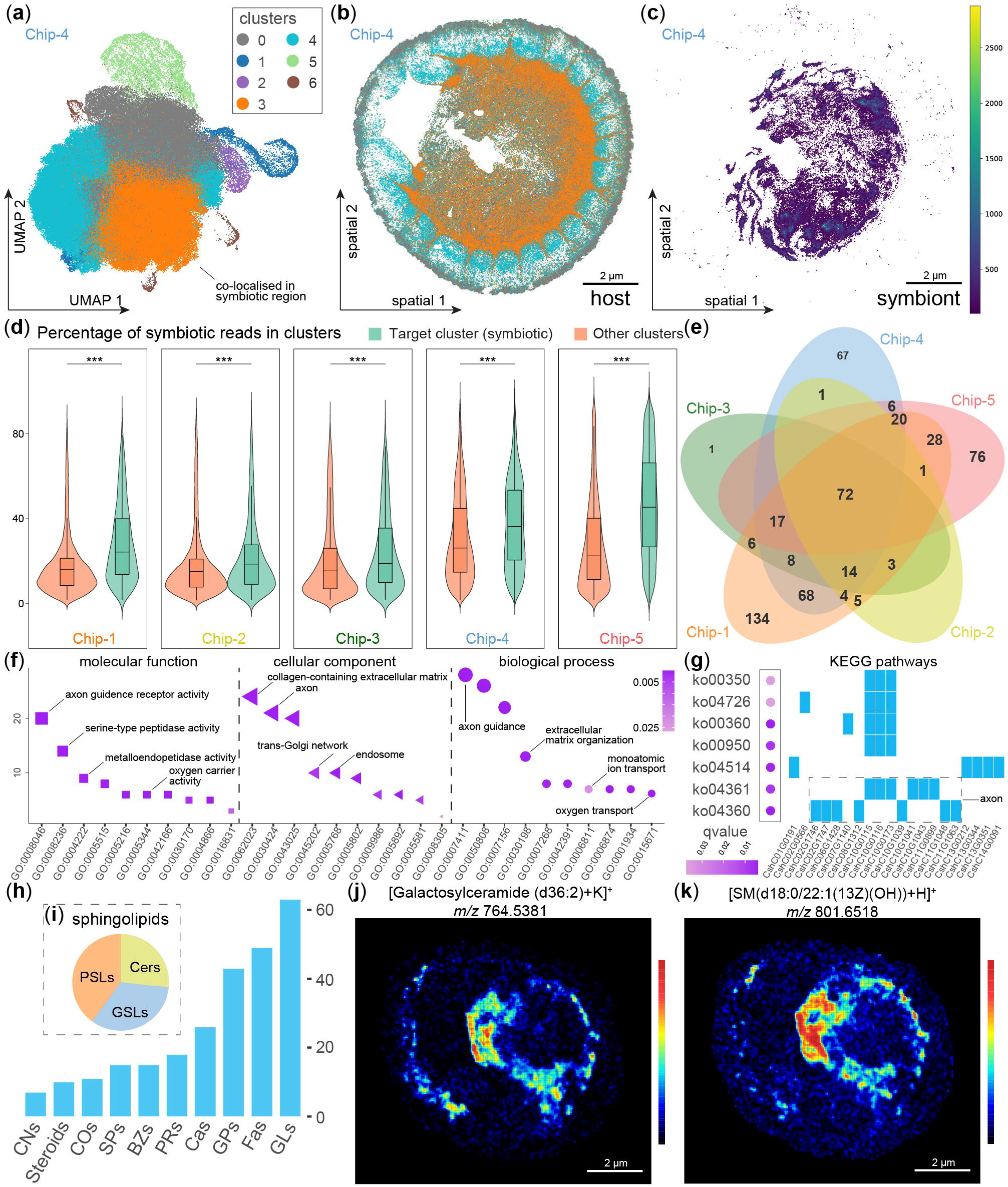
**a**, Uniform manifold approximation and projection (UMAP) of clusters from aggregated bins (host) in chip-4, with the information of all the five chips shown in Supplememtary files Table S7. **b**, Spatial representation of clusters (host) in chip-4. **c**, The spatial representation of symbiotic reads in chip-4. **d**, The percentage of symbiotic reads in symbiotic clusters and the others. **e**, Venny plot of marker genes in symbiotic clusters from five independent analysis, with candidate genes labelled in bold if the identification in more than 2 results. **f**, Functional category of candidate genes based on the gene ontology. **g**, Functional category of candidate genes based on the KEGG pathway. **h**, Histogram plot of identified metabolites in DESI-MSI at the class level. **i**, Pie plot of three categories of 15 identified metabolites affiliated to sphingolipids. **j**, Spatial pattern of galactosylceramide (m/z 764.5381). **k**, Spatial pattern of sphingomyelin (m/z 764.5381). GLs: glycerolipids, Fas: fatty acyls, GPs: glycerophospholipids, Cas: carboxylic acids and derivatives, PRs: prenol lipids; BZs: benzene and substituted derivatives, SMs: sphingolipids, Cos: organooxygen compounds, Steroids: steroids and steroid derivatives, CNs: organonitrogen compounds.

Spatial transcriptomics revealed an enriched pattern of proteolysis-related genes in the digestive gland, including serine-type peptidase and metallopeptidase activities (Fig. 3f). These could aid the digestion of symbiont proteins as a source of amino acids^45^, which would complement the deficiency of the host’s biosynthesis capacity (Extended Data Fig. 6 and Supplementary files Table S13). We did not observe lysosomes in dracospherae, but densely packed lysosomes (~200 nm diameter) were present in the adjacent cytoplasm (Fig. 2 and Extended Fig. 4b-c), and we also found evidence of hexosaminidase (enzyme in lysosome, Supplementary files Table S15) in the symbiotic region. The multi-membranes of the dracosphera prevent lysosomes from entering the dracosphera, and the enormous size of the dracosphera also hinders direct digestion via traditional phagocytosis, unlike other chemosymbiotic holobionts. Therefore, we hypothesise that the dracosphera may ‘burst’ at some point to release the symbionts into the cytoplasm – which can then be digested by the lysosomes (Fig. 4).

**Fig. 4.**
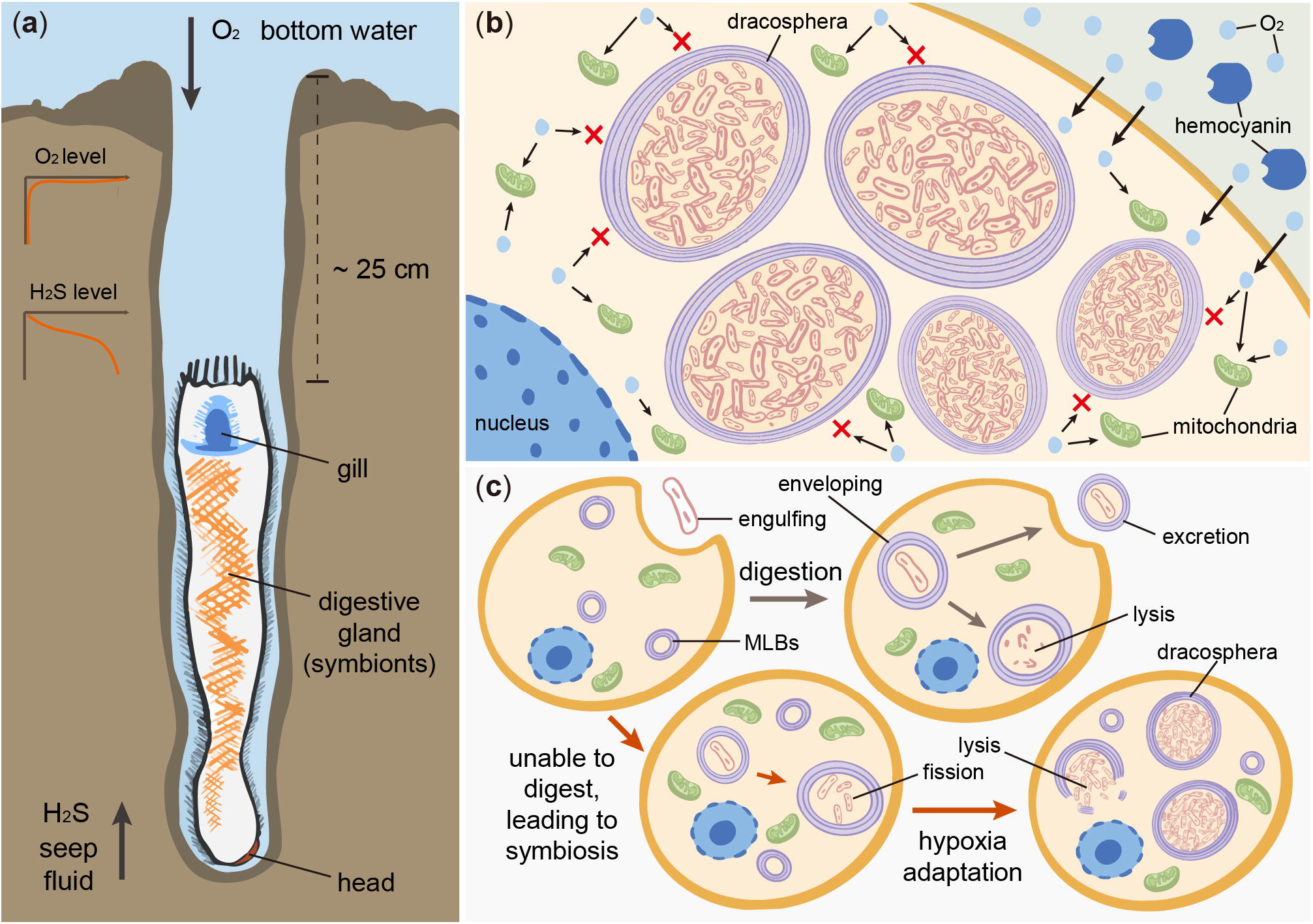
Schematic illustration of the chemosymbiosis in *Chaetoderma shenloong* using the specialised organelle, dracosphera. **a**, Anatomy and life position of *C. shenloong* buried deep in anoxic seep sediment to take in hydrogen sulfide from the posterior gill. **b**, Concept illustration for the inferred function of the dracosphera, where oxygen entering the host bateriocyte cannot diffuse through to the symbionts due to the multilamellar wall of the dracosphera and is instead taken up by host mitochondria. **c**, Proposed evolutionary pathway for dracosphera. Green arrows indicate the likely original function of the multilamellar bodies (MLBs) where they are secreted to envelop and digest or excrete bacteria engulfed by the animal cell. Red arrows indicate the pathway where a sulfur-oxidising bacteria was able to resist lysis inside MLBs, and instead multiplying inside them, eventually leading to a symbiosis with host. With further adaptation to hypoxia, the membranes increased in layering to form the dracosphera to allow the host cell to fully utilise all available oxygen.

### Potential function of the dracosphera

Why would *C. shenloong* compartmentalise its symbionts into dracospherae when other chemosymbiotic organisms manage without such costly organelles^2^? One explanation lies in its adaptation to deep, anoxic, reducing environments. The habitat of *C. shenloong* is hypoxic, a typical challenge for vent and seep holobionts that must adapt to obtain both reducing chemicals and oxygen^38^. However, given its burrowing depth, *C. shenloong* may represent an extreme case even among chemosymbioses. Hypoxic and anoxic conditions can drive evolutionary innovations, such as the transformation of mitochondria into hydrogenosomes in loriciferans inhabiting permanently anoxic sediments up to 15 cm below the surface^5,39^. While *C. shenloong* likely extends its posterior gill to the surface for oxygenation, it also needs to burrow deep to take in hydrogen sulphide, which is only available at greater depths (Extended Data Fig. 1b). Consequently, *C. shenloong* may need to endure prolonged periods of anoxia.

Ultrastructure of the dracosphera superficially resembles other multilamellar elements known in nature, such as myelin sheaths around neural axons that both function as insulators and produce ATP by oxidative phosphorylation, as well as thylakoid membranes in cyanobacteria which are sites of photosynthesis^40^. A key function of these multilamellar elements is the absorption and release of gases, especially oxygen and carbon dioxide. Given the *C. shenloong* endosymbiont is facultatively anaerobic and does not require oxygen, it is plausible that the host mollusc would benefit from preventing the symbiont from contacting oxygen; thereby retaining all available oxygen for use by the animal under hypoxic conditions. Furthermore, the *C. shenloong* symbiont may function more optimally under hypoxic conditions, though the close relatives of the symbiont are not obligate anaerobes^32^. The multilamellar wall of dracospherae possibly functions as a barrier of oxygen diffusion, and at the same time conversely delivers carbon dioxide (Fig. 4b). The diazotroph *Frankia* symbiotic with angiosperm plants is known to use multilamellar external vesicles to defend against high oxygen concentrations since nitrogenase is oxygen-sensitive^3^. Many termites house anaerobic symbionts in their hindgut to break down lignin and cellulose, which the host accommodates by constructing an oxygen gradient inside the body^41^. It has also been speculated that many such structures have convergently evolved for energy production^40^.

The structural similarity between myelin sheath and dracosphera is corroborated by the high expression of genes known to be linked with axon activities and the nerve system such as fasciclin 2, semaphorins, and acetylcholine receptors (Supplementary files Table S15) in the symbiotic organ. Furthermore, 15 sphingolipids (Fig. 3h-i) were enriched in the digestive gland, identified from spatial metabolomics powered by desorption electrospray ionization mass spectrometry imaging (DESI-MSI, resolution: 50 μm).

Among them, galactosylceramide (Galc) and sphingomyelin (SM) (Fig. 3j-k and Extended Data Fig. 8) are the major structural components in mammal nerve systems^46^. Ceramides, the precursors and products of these genes, also exhibited a similar pattern indicating the active transformation among them (Extended Data Fig. 7 and Supplementary files Table S15). Comparing the ambient cytosolic fluid, the lipid-rich profiles in the dracospherae were further supported by Raman spectrums (Extended Data Fig. 9). Six hemocyanins were highly expressed in the digestive gland (oxygen carrier activity in Fig. 3f and Supplementary files Table S15), which would be responsible for oxygen transportation.

The dracosphera could have arisen because of how the symbiotic relationship evolved. In the social amoeba *Dictyostelium discoideum*, facultative symbiosis with the bacteria *Burkholderia* is a by-product of the amoeba packaging bacterial cells with multilamellar membranes for digestion^42^. *Burkholderia* can resist digestion because the multilamellar envelope helps it tolerate harsher conditions^43^. We speculate that the symbiosis between *C. shenloong* and its symbiont may have originated in a comparable way (Fig. 4c), with the dracosphera protecting the bacteria and the bacteria providing energy in return. As multilamellar bodies can now be synthesised using artificial cell techniques^44^, this mode of establishing symbiosis could be tested in the future via experimental enveloping of bacterial lineages. Though chemosynthesis is ubiquitous in marine realms, the dracosphera is the first organelle dedicated to it and presents a new opportunity to understand organellegenesis, pending future studies on its origin and machinery.

Our discovery of a remarkable adaptation from one of the most poorly understood animal groups highlight how little we know about the potential of marine invertebrates, especially infaunal deep-sea taxa^47^. During this study, we serendipitously collected an individual of *C. shenloong* from a seep site near a mud volcano (‘MV3’) off Kyushu in Japan. This extends the range of this species from two seeps in the South China Sea (Haima and Jiaolong Ridge^22,23^) to Japan. Despite its enormous size and unusual ecology, *C. shenloong* has remained undiscovered across its wide range until recently. The dracosphera enables aplacophorans, and other animals, to live much deeper in anoxic mud than previously thought. Given the largest known living caudofoveate (*C. felderi* reaching 40 cm length) resides in cold seeps in the Gulf of Mexico^22,48^, chemosymbiotic aplacophorans may be much more common than we realise and contribute significantly to primary production at seeps and the deep ocean as a whole^1^. How many more such evolutionary marvels with potential significance in geochemical cycling remain hidden and unaccounted for, in the ‘last frontier of mankind’, which faces imminent risks from human activities in the deep sea?

## Acknowledgements

This work was supported by the National Key Research and Development Program of China (2024YFC2816100), the Science and Technology Innovation Project of Laoshan Laboratory (LSKJ202203104), Natural Science Foundation of Shandong Province (ZR2023JQ014), the grant from Natural Science of China (HJRC2023001), the PI project (2021HJ01) from the Southern Marine Science and Engineering Guangdong Laboratory (Guangzhou), two Fundamental Research Funds for the Central Universities (202172002 and 202241002), and the Collaborative Research Fund (C2013-22GF) from the Hong Kong SAR Government. We thank the captain and crew of R/Vs *Xiangyanghong 01* and *Hakuho-Maru*, as well as ROV *Pioneer*, during cruises XYH01-2022-06 and KH-23-4 Leg 2 (PI: Akira Ijiri, Kobe University). Hiromi K. Watanabe (JAMSTEC) kindly collected and provided a specimen of *Chaetoderma shenloong* from off Kagoshima, Japan for this study.

## Author contributions

Conceptualisation: J.S. Methodology: Y.L., C.C., X.L., M.L., H.W., X.H., S.L., and J.S. Morphological investigation: C.C., and X.L. Genome and spatial transcriptomics: Y.L. Spatial metabolomics: Y.L. H.Z. Bioinformatics: Y.L. Staining: M.L., X.L., and H.W. Electron microscopy: Y.L. Visualisation: Y.L. and C.C. Funding acquisition: J.S., J.-W.Q., and P.-Y.Q. Project administration: J.S. Supervision: J.S. Writing – original draft: Y.L. and C.C. Writing – review & editing: X.L., M.L., H.W., X.H., S.L., J.-W.Q., P.-Y.Q., and J.S.

## Competing interests

We declare that we have no competing interests.

## Data and materials availability

The raw sequencing reads in this work have been deposited in NCBI bioproject, with the genome assembly (i.e., HiFi, Hi-C, RNA-seq) in PRJNA1149698, genome resequecing reads in PRJNA1206971, spatial transcriptomics in PRJNA1206974. The assembled genome and its gene feature files are available at Figshare (https://figshare.com/s/ef2275353b0cd547f298), including the host and symbiont. The commands in this study, including genome assembly, gene model prediction, SNP callings, and functional enrichment spatial transcriptomics, are available at GitHub (https://github.com/yligy/Csh).

---

**Extended Data Fig. 1 a**, Known distribution of *Chaetoderma shenloong*. **b**, Push-corer as recovered from the seafloor, with live *C. shenloong* individuals. **c**, Close-up of specimens showing the head-down position of *C. shenloong* in the sediment. Yellow arrows indicate the oral shield and white ones indicate the neck. Concentration vs sediment depth of **d**, oxygen, **e**, hydrogen sulphide, and **g**, nitrate within the same push core.

**Extended Data Fig. 2 a**, The binding range of 16S rRNA of the sFISH probes. **b**, Signal of EUB338 probe targeting eubacteria in cyan. c, Signal of specific probes targeting *Chaetoderma shenloong* symbiont in red. **d**, Signal of DAPI targeting nucleic acids in blue, with strong signal for host nucleus and lower signal for symbiont. **e**, Signal of CellMask targeting cell membrane in green. **f**, Merged view of eubacteria (b), symbiont (c), DAPI (d), and membrane (e). **g**, Colocalization of eubacterial signal and specific symbiont signal, for the line shown in part f. **h**, Line plot showing the intensity of signals along the line shown in part f. **j**, Example image showing how the diameter of dracospherae was measured using ImageJ. **K**, Violin plot of the dracosphera diameters in the anterior, middle, and posterior parts of the animal.

**Extended Data Fig. 3 a**, Stable isotope compositions (δ^13^C and δ^15^N) of macrofauna collected from Haima seep. Most of the data (labelled as triangle) were published in a former study and re-plotted (see fig. 5a in Li et al., 2023) except for *Chaetoderma shenloong*. The blue and light red indicate that the tissues harbours methane-oxidizing bacteria (MOB) and sulphur-oxidising bacteria (SOB) symbiont, respectively. **b**, Genome evaluation of *C. shenloong* using GenomeScope. **c**, Heatmap plot showing the Hi-C contact intensity during 3D-DNA scaffolding. **d**, Circos plot of the pseudo-chromosome level genome of *C. shenloong*. The three sub-rings represent the GC content, gene numbers, and repeat contents in 1 Mb window size, respectively from the outside inwards. **e**, Phylogenetic position of *C. shenloong* with selected molluscs and four outgroup taxa with high-quality genome data (IQ-TREE2, best-fitting model with partition)

**Extended Data Fig. 4 TEM micrographs showing ultrastructure of the digestive gland and dracospherae. a**, A dracosphera wrapped by a membranous structure. **b**, Several dracosphera densely packed together. **c**, cytoplasmic structure containing lysosomes, with a magnified viewin **d**.

**Extended Data Fig. 5 Genomic information of the sulfur-oxidising *Thiodubiliella* symbiont of *C. shenloong*. a**, Circos plot of the newly assembled and complete symbiont genome. The four ring plots inside represent feature in forward, feature in reverse, GC level, and GC skew, from the outside inwards. **b**, Genomic comparisons of *Thiodubiliella* symbionts, with genomic size and the number of coding sequences (CDS). **c**, Histogram plot of SNP density in the symbiont, none of SNP was identified from the individual used for genome sequencing. **d**, Histogram showing the coverage (per nucleus) of mitochondria and symbiont in samples using genome resequencing. The value was calculated as the coverage of them divided by the 50% haploid coverage of host.

**Extended Data Fig. 6 Overview of metabolic pathways in the *Chaetoderma shenloong* symbiont**. Solid arrows represent the presence of genes or enzymes in the symbiont, whereas the dashed arrows indicate absence. Solid black arrows indicate the enzymes or genes that were identified as highly expressed (top 300 TPMs), others are indicated in light grey. Orange arrows indicate that the absence of a gene in the symbiont is compensated by that of the host. Animo acids shown in solid black font have complete biosynthesis pathway in the symbiont, and others are shown in light grey.

**Extended Data Fig. 7 Symbiont signal in FISH, symbiotic reads in spatial transcriptomics, symbiotic cluster, and UMAP distribution**. The row names indicate the different chips (sections) of *Chaetoderma shenloong*, with the detail shown in Supplementary files Table S7. The column names indicate the symbiotic signal in FISH, distribution of symbiotic reads in the spatial transcriptomics, distribution of symbiotic cluster, and UMAP distribution.

**Extended Data Fig. 8 Spatial mapping of DESI-MSI in sections containing the symbiotic digestive gland**. The row names indicate the different sections of *Chaetoderma shenloong*, No. 2 is around the neck, No. 4 is middle of the body (trunk), No. 8-1 and No. 8-2 for the posterior trunk, anterior of the pericardium. The column names indicate the symbiotic signal in FISH, the merged signal from DAPI, symbiont, and membrane, and three sphingolipids.

**Extended Data Fig. 9 Raman spectra under microscopy in sections of the digestive gland. a**, Images showing the location of the Raman spectra. **b**, Raman spectra of 8 locations. **c**, Two typical spectra showing the characteristics of peaks (details shown in Supplementary file Table S20).

## Notes

### Competing Interest Statement

The authors have declared no competing interest.

