## Supplementary Figures for "A multilamellar organelle for chemosymbiosis in an aplacophoran mollusc adapted to anoxic cold seep sediment"

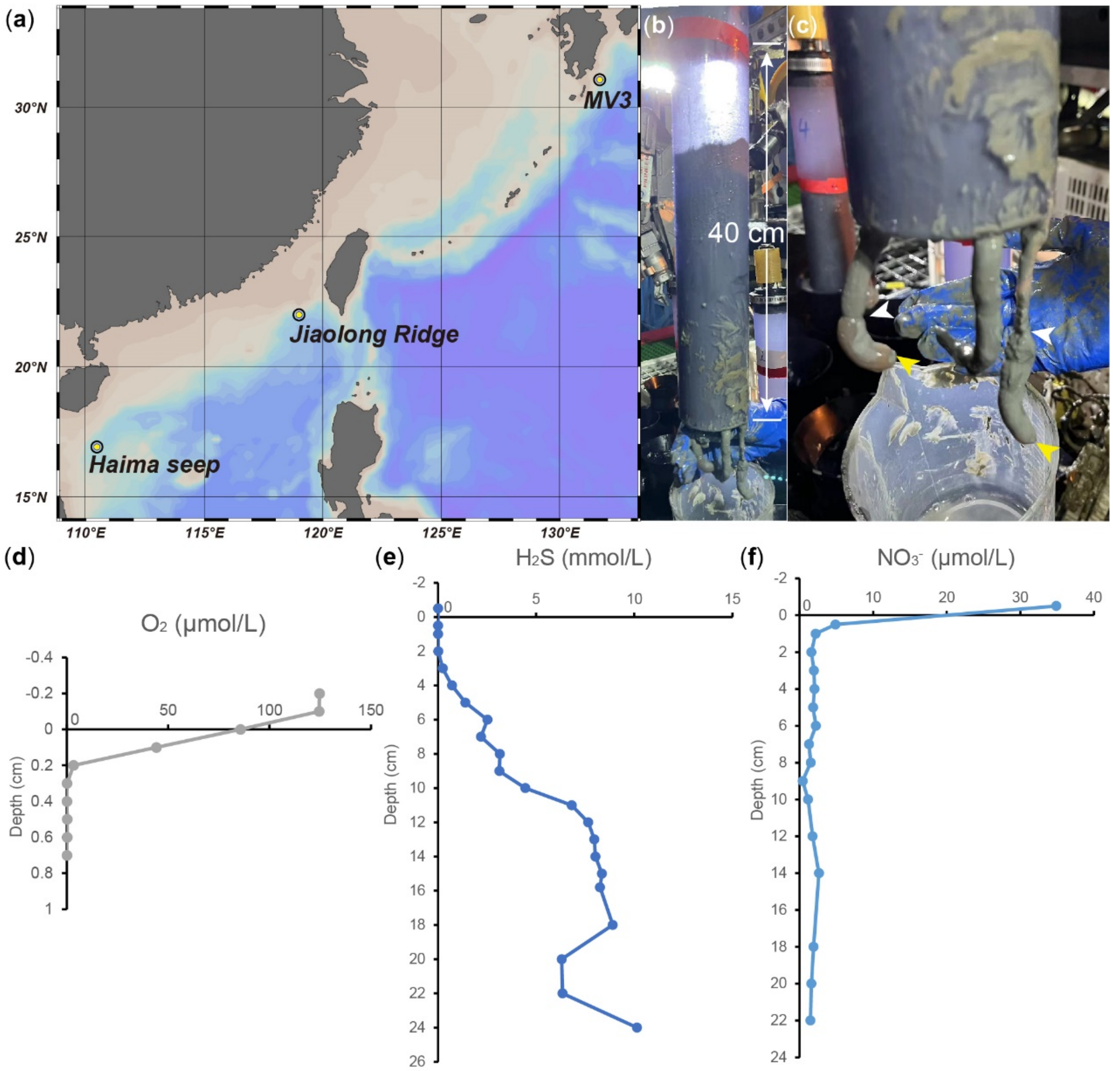


Extend Data Fig. 1

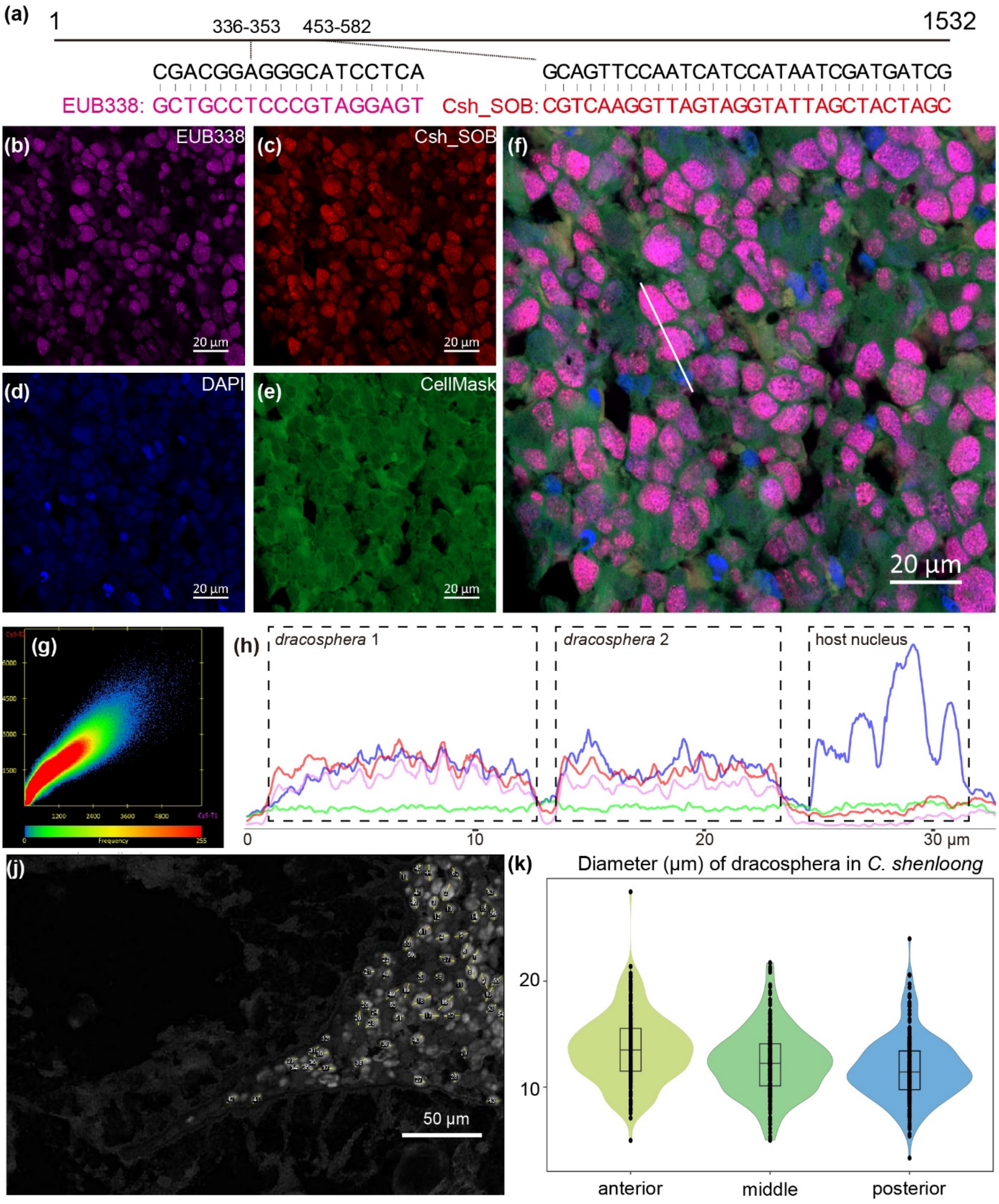


Extend Data Fig. 2

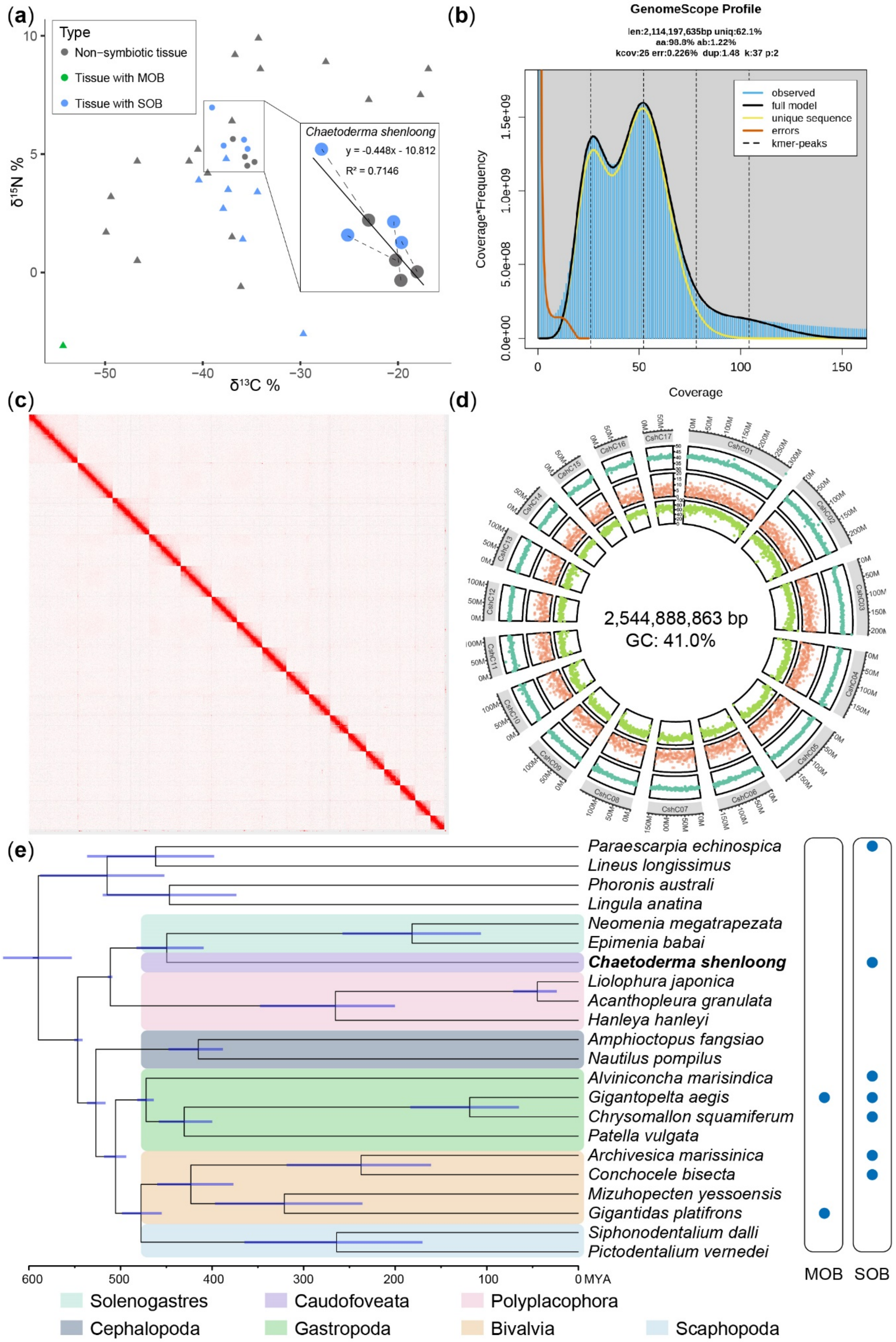


Extend Data Fig. 3

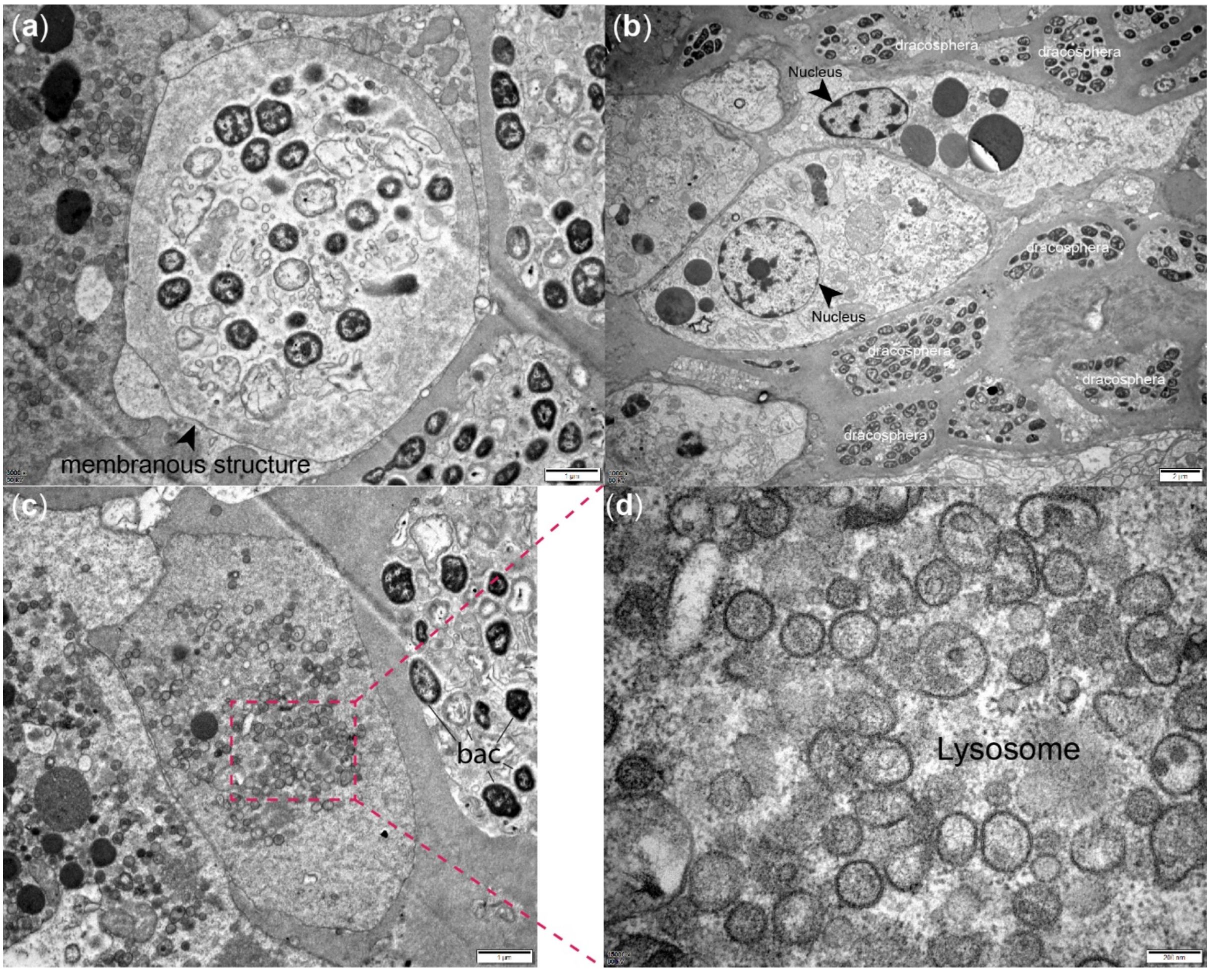


Extend Data Fig. 4

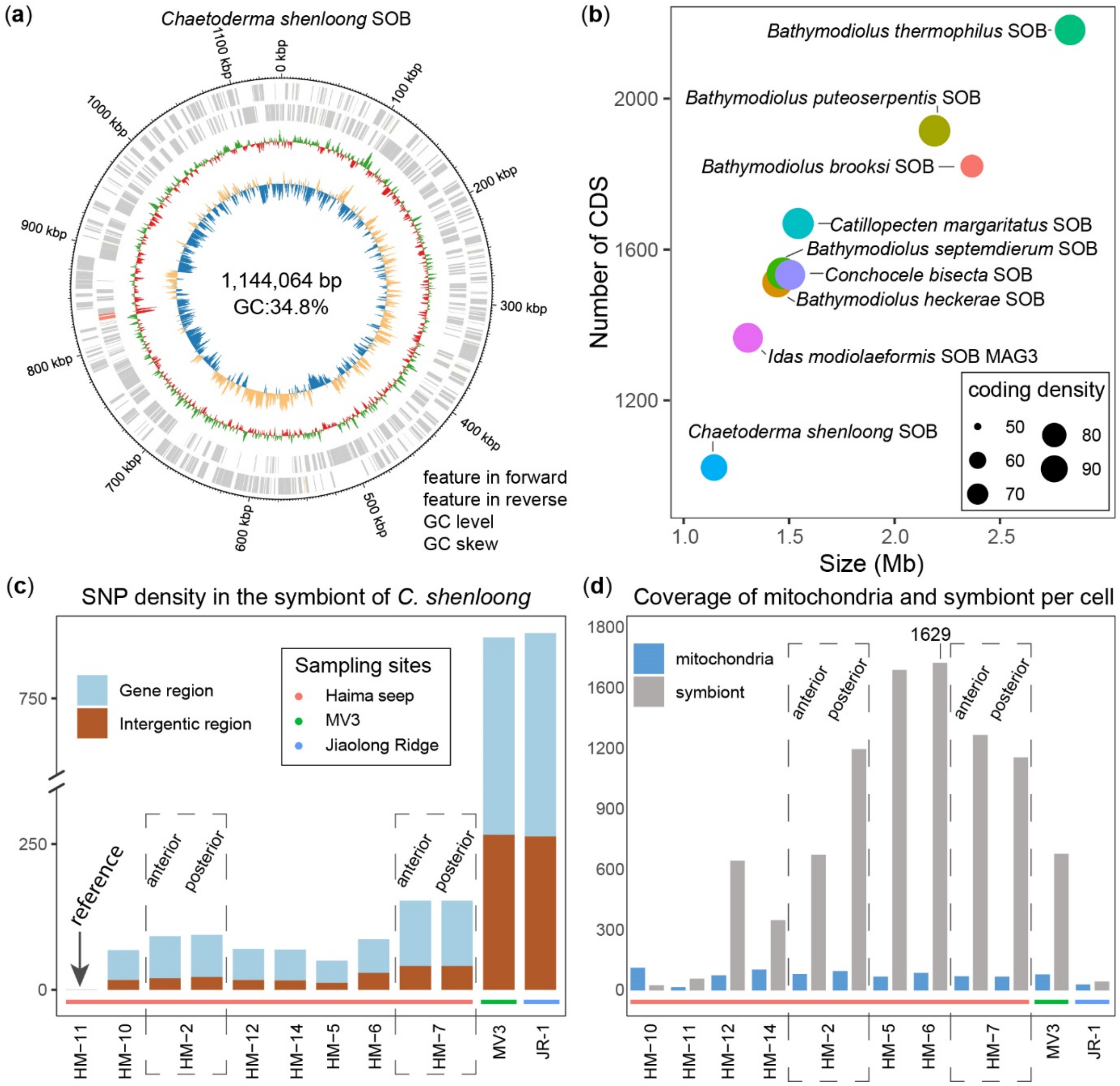


Extend Data Fig. 5

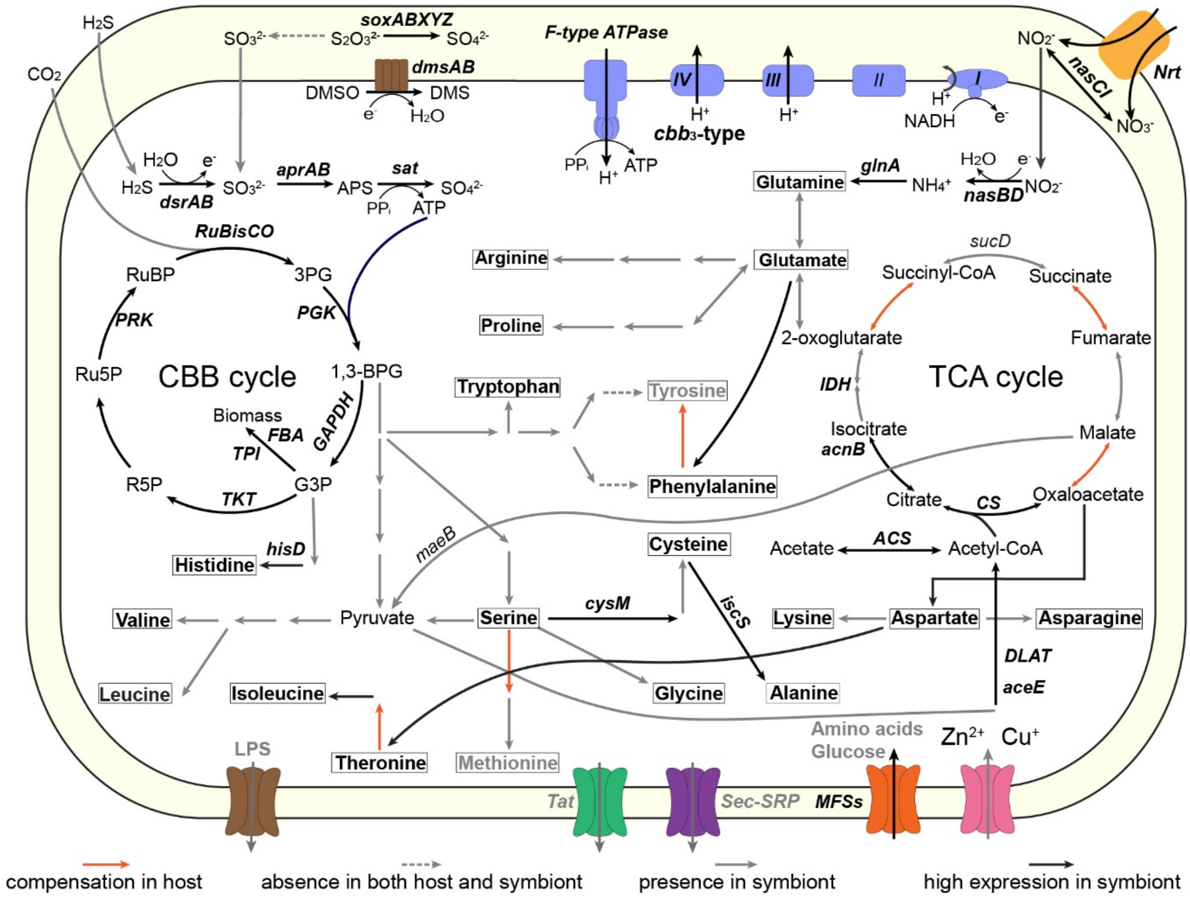


Extend Data Fig. 6

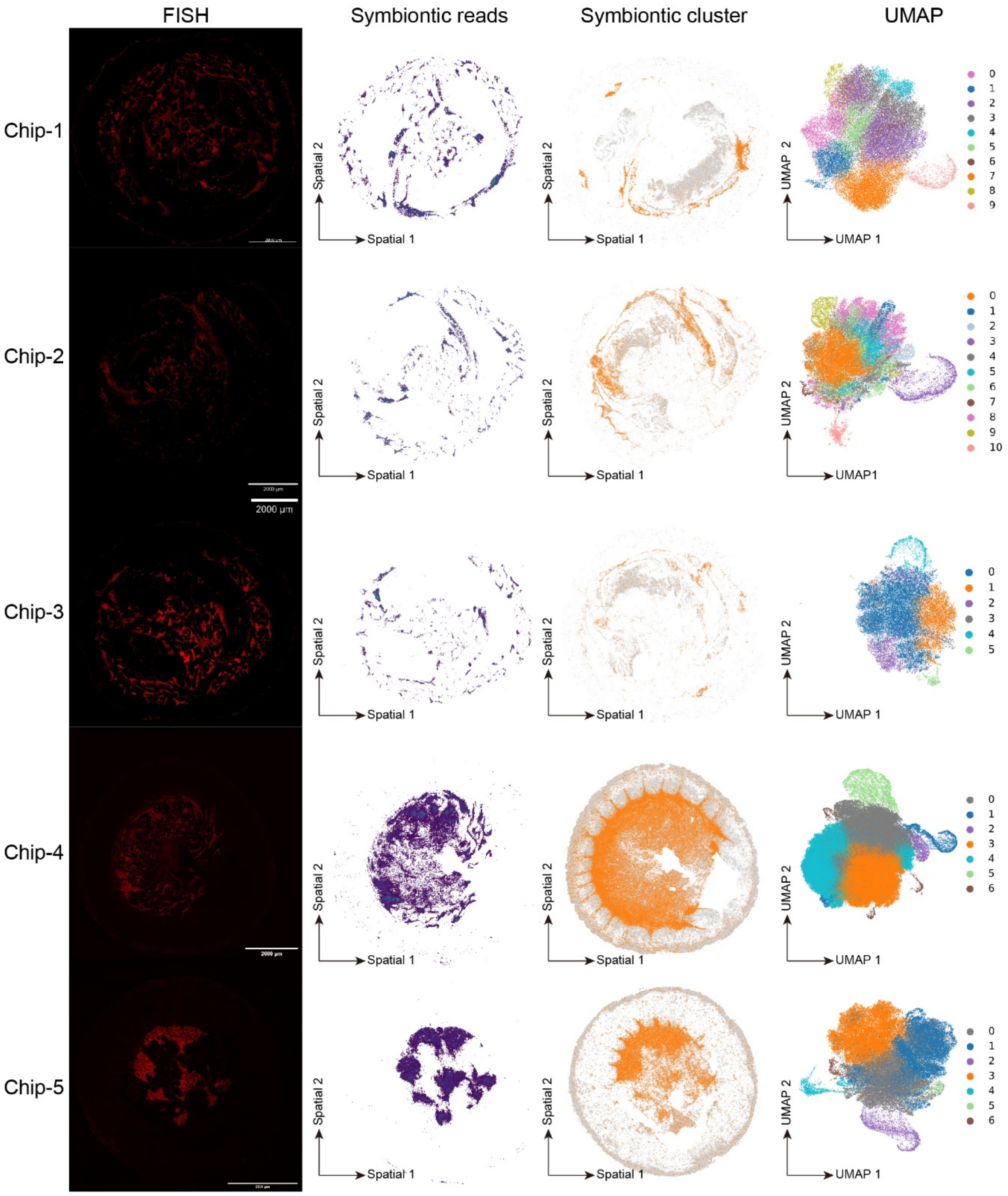


Extend Data Fig. 7

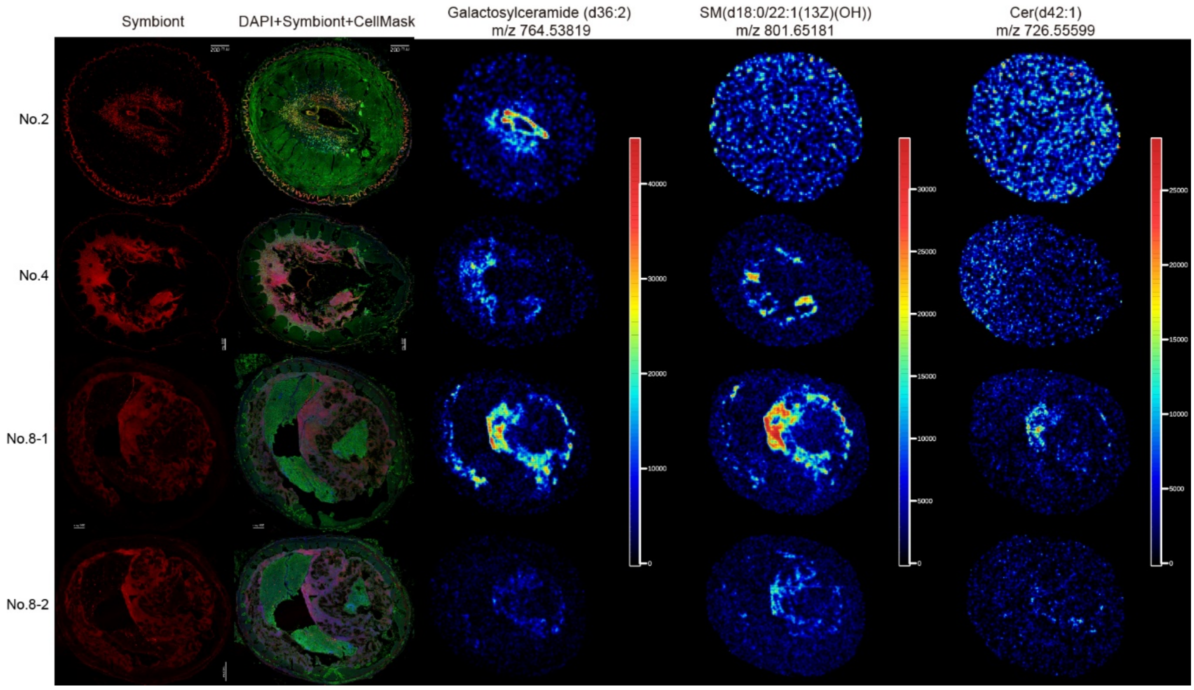


Extend Data Fig. 8

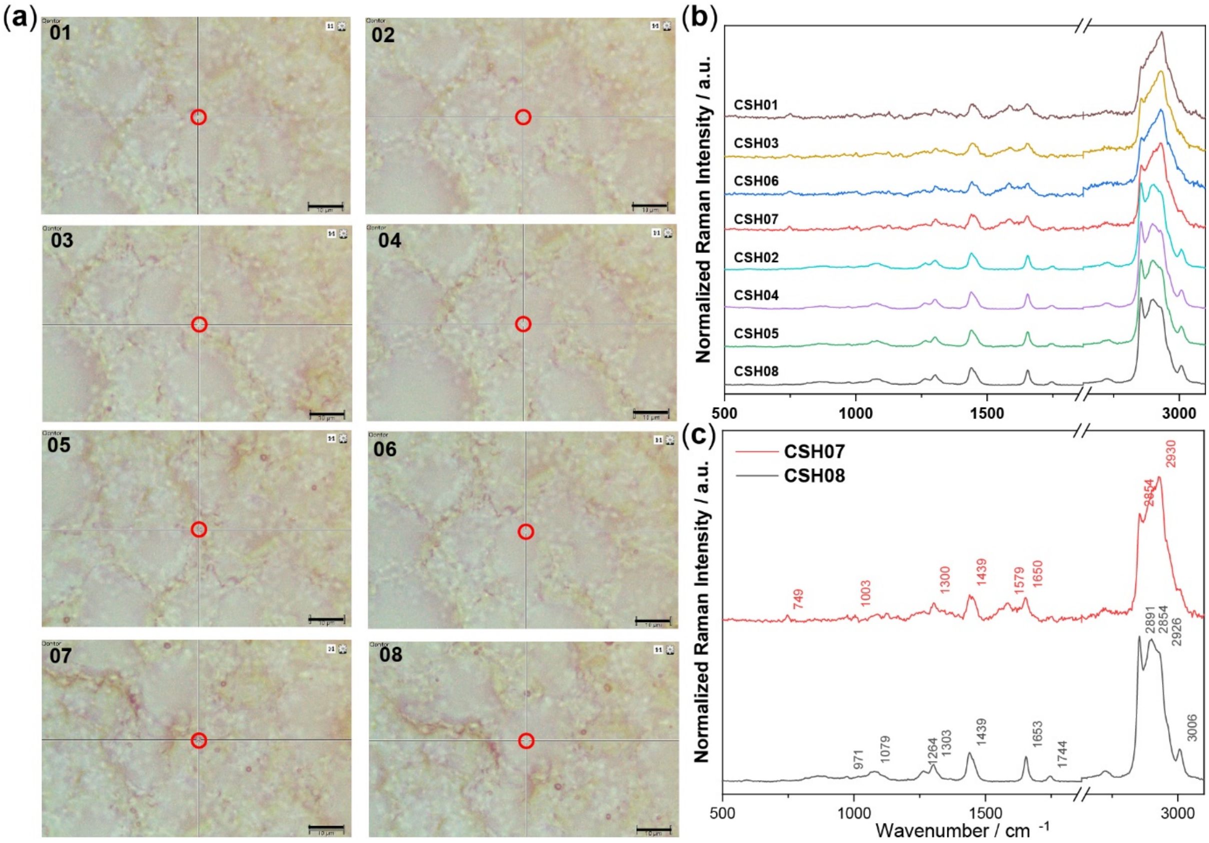


Extend Data Fig. 9
