## Supplementary Methods for "A multilamellar organelle for chemosymbiosis in an aplacophoran mollusc adapted to anoxic cold seep sediment"

*Sample collection and preservation*

Eight individuals of *Chaetoderma shenloong* were collected from Haima methane seep in the South China Sea using a 50 cm long push-core mounted on the remotely operated vehicle (ROV) *Pioneer* on the research vessel (R/V) *Xiangyanghong 01* of the First Institute of Oceanography, Ministry of Natural Resources (MNR), China, September 2022. One other individual was collected from a cold seep associated with the MV3 mud volcano (Mitsutome et al. 2023) off Tanegashima, Japan by an Okean grab on-board R/V *Hakuho-Maru* cruise KH-23-4 Leg 2 (PI: Akira Ijiri, Kobe University). For detailed collecting data see Supplementary file Table S1.

Samples were preserved in multiple ways after recovery on-board the research vessels, using the following methods: 1) flash-frozen in liquid nitrogen in 15ml tubes and then stored at -80°C; 2) specimens were fixed in RNAlater (ThermoFisher, USA) and then stored at -80°C; 3) fixed in 4% paraformaldehyde fix solution (PFA) overnight, then substituted by 100% methanol and stored at -20°C; 4) fixed and stored in 2.5% glutaraldehyde solution at 4°C; 5) fixed and stored in 95% ethanol at room temperature. The usage of these samples for different analyses is shown in Supplementary file Table S5.

*μ-CT scanning and 3D reconstruction*

The specimen fixed by PFA and transferred to methanol was subjected to a rehydration series with a decreasing concentration of methanol to Phosphate Buffered Saline (PBS) solution (90%, 80%, 70%, 60%, 40%, 20%) for 1 hour each, and then placed into pure PBS for 24 hours. The spicules were first roughly removed with scalpels and needles, and then placed in 0.1 M HCl solution until complete decalcification as indicated by a lack of emerging bubbles. The specimen was then stained with 3% Lugol’s solution for two days.

A SkyScan 1276 μ-CT scanner (Bruker, Belgium) was used to scan the specimen, placed in a 50 ml centrifuge tube filled with water. The scan was conducted at 55 kV and 200 µA, at a pixel resolution of 6.54 µm. The software NRecon (Bruker, Belgium) was used to obtain the 2D sections based on the scanned X-ray images. These sections were processed with CTAn and DataViewer software (Bruker, Belgium). The specialist software Amira v2022.2 (Thermo Fisher Scientific) was used for 3D reconstruction of the internal organ, by manually highlighting the organs in each section and then generating the 3D surface.

*Scanning electron microscopy*

A specimen preserved in 2.5% glutaraldehyde solution was cut into small pieces (about 1~2 mm thickness) and then washed in 0.1M PBS for 15 min, repeated three times. The specimen was cut to small pieces 1-2 mm in thickness, and then critical-point dried. The dry specimens were sputter-coated with gold (Au), and then imaged using a VEGA3 SEM (TESCAN) at 10.0 kV.

*Transmission electron microscopy*

Tissue from a specimen fixed in 2.5% glutaraldehyde solution was cut and washed in the same manner as for SEM observations. Subsequently, the tissue was fixed in 1% osmium tetroxide (OsO^­^_4_) for 1 hour and then washed in 0.1M PBS for 15 min, for three times. The specimen was dehydrated in an increasing methanol series (50, 70, 90, and 100%, for three times each) for 15 min each and embedded in resin (Epon 812). Thin-sections at 70 nm thickness was cut using a Reichert ULTRACUT Ultrathin slicer (Leica, Austria). The thin sections were stained using the uranyl acetate and lead citrate double staining method (Bernhard 1969), with a uranyl acetate and lead citrate staining for 15 min each. Images were captured using a JEM 1200-EX (JEOL, Japan) TEM at an accelerating voltage of 80 kV.

*Focused-ion beam scanning electron microscopy*

We used a focused-ion beam scanning electron microscope (FIB-SEM) to construct the 3D structure of the dracosphera. The specimen was stained using the following procedure: washed in 0.1M pH 7.2 phosphate buffer (PB) for 3 min for three times; stained in 0.1M PB containing 1.5% ferrocyanide and 2% OsO^­^_4­_ for 1 hour under dark environment and on the ice then washed in double-stilled water for 3 min for five times; stained again in 1% thiocarbohydrazide (TCH) solution at room temperature for 20 min followed by washing in double-stilled water for 3 min for five times; stained in 2% OsO^­^_4_ in water for 30 min under dark environment and on the ice then washed in double-stilled water for 3 min for five times; stained in 1-2% uranyl acetate in water at room temperature for 30 min then washed in double-stilled water for 3 min for four times and again at 60 °C oven for 3 min; and finally in Walton’s Lead (39.9 mg L-aspartic acid and 66 mg lead nitrate in 10 ml double-stilled water, pH=5.5) at 60 °C oven for 3 min and then washed in 60 °C double-stilled water for 3 min for five times.

The stained specimen was dehydrated specimen in an increasing serial ethanol solution (30%, 50%, 70%, 80%, 90%, 95%), twice in each solution for 5 min, then twice in 100% ethanol for 10 min. Resin embedding was done in the following steps: 5 mins in 1:1 ethanol/acetone solution, 1 hour in resin/acetone solution (1:2) at room temperature followed by 1 hour in 1:1 resin/acetone solution and then 1 hour in 2:1 resin/acetone solution (2:1), and finally in pure resin at room temperature for 1 hour, and repeated three times. The resin-embedded specimen was hardened in a 60°C oven for 2 days. A Crossbeam350 (ZEISS, Germany) FIB-SEM was used to image the specimen at the resolution of 10 nm.

*Metagenome sequencing and assembly*

Genomic DNA was extracted from the symbiotic digestive gland using the SDS method and then randomly broken into about 350bp fragments. After library construction, it was sequenced with the paired-end 150bp mode in an Illumina Novaseq 6000 platform. The raw reads were trimmed in trimmomatic version 0.39 (Bolger et al. 2014) with the following settings (TruSeq3-PE-2.fa:2:30:10:8:true SLIDINGWINDOW:5:20 LEADING:3 TRAILING:3 MINLEN:36). MEGAHIT version 1.2.9 (Li et al. 2015) was used in metagenome assembly with the default settings. MaxBin2 verion 2.2.7 (Wu et al. 2015) was used to retrieve the symbiont genome and the fragmented host genome. We also downloaded and reanalysed reads from a previous work on *Chaetoderma shenloong* (Wang et al. 2024).

*Genome sequencing and assembly*

Muscle and gill tissues from a flash-frozen individual were used in high-molecular weight DNA extraction using the SDS method. The quality-checked DNA was sent to Novogene Co. Ltd (Tianjin, China) for library construction and sequencing. High fidelity (HiFi) sequencing was adopted in the genome assembly, with the platform and outputs shown in Supplementary Table S6. Meanwhile, a part of tissue was processed in high-throughput chromosome conformation capture (Hi-C) library with the restriction emzyme MBoI as the guideline, and the PE-150 reads were output from an Illumina Novaseq 6000 platform.

Reads were extracted from .bam file using extracthifi version 1.0.0 and then the adapters were checked using HiFiAdapterFilt version 2.0. Before assembly, minimap2 version 2.24-r1122 (Li 2018) was employed in mapping HiFi reads to the draft genome of the symbiont (from the NGS-based metagenome). Therefore, HiFi reads were divided into the symbiont reads and the symbiont-free reads. The symbiont HiFi reads were subjected to hifiasm_meta version 0.3-r063.2 (Feng et al. 2022), which successfully assembled the complete genome of symbiont. The symbiont-free reads were subjected to hifiasm version 0.16.1-r375 (Cheng et al. 2021) for genome assembly with the default settings. The duplicated and contaminated contigs in the primary assembly were identified using minimap2, blastn (Camacho et al. 2009), Purge_Dups version 1.2.5 (Guan et al. 2020), TaxonKit version 0.15.1 (Shen and Ren 2021), diamond blastp version 2.0.15 (Buchfink et al. 2021), and MEGAN version 6.21.7 (Gautam et al. 2022), which resulted in the non-redundant and decontaminated contigs of *C. shenloong*.

To anchor these contigs to the superscaffold level or pseudo-chromosome level, the Hi-C reads were processed in HiC-Pro version 3.1.0 (Servant et al. 2015) to remove the unlinked reads. Then, they were subjected to Juicer version 1.6 (Durand et al. 2016) for the output of merged_nodups.txt and then the contigs were subjected to the 3D de novo assembly (3D-DNA) pipeline version 201008 (Dudchenko et al. 2017). Juicebox Assembly Tools version 2.17.00 (Dudchenko et al. 2018) were employed to generate and visualize a Hi-C contact map and then the mis-assembly or mis-scaffolding were manually corrected. Lastly, the reviewed links were subjected to 3D-DNA for constructing the pseudo-chromosome level genomes. The completeness of genome, transcriptome, and the proteome (see below) were evaluated using Benchmarking Universal Single-Copy Orthologs (BUSCO) version 5.2.2 (Manni et al. 2021) against the metazoan odb10.

*Repeat region and gene model prediction of the host genome*

The general pipeline was slightly modified from a published work (Sigwart et al. 2024). Briefly, the soft-masked genome and the corresponding repeat level were called using RepeatModeler v2.0.3 (Flynn et al. 2020) and RepeatMasker v4.1.2-p1 (Smit et al. 2015). To obtain the transcriptional profile, the RNALater-preserved sample was dissected into different organs (gill, muscle, digestive gland, pericardium, gonad, and epidermis) and then used in RNA-seq (details shown in Table SX). In trimming metagenomic reads, the low-quality or adapter-contaminated RNA-seqs were removed using trimmomatic. These reads were then aligned to the soft-masked genome using aligner STAR version 2.7.10a (Dobin et al. 2012), which generated .bam files according to mapping regions. BRAKE2 version 2.1.6 (Brůna et al. 2021), implemented with Augustus version 3.4.0 and GeneMark version version 3.67_lic for the ab inito gene prediction, was employed in *ab-initio* prediction (“augustus.gff3” and “genemark.gff3”). The protein-coding regions were also predicted via homolog alignment, with proteins from 28 high-quality genomes in Metazoa aligned to the soft-masked genome using miniport version 0.5-r179 (Li 2023) (“proteins.gff3”). Trinity version 2.13.2 was adopted in assembling transcripts with both de novo and genome-guided modes, whose outputs were aligned to genome using PASA version 2.5.2 (Haas et al. 2008). Meanwhile, Stringtie v2.1.1 (Pertea et al. 2015) was also used in predicting the expressed regions in genome, which was used in Evidencemodeler version 2.1.0 (Haas et al. 2008) to integrate the above three predictions into a comprehensive profile of genes, and the weights of evidence were AUGUSTUS for 2, GeneMark for 2, PASA for 10, Stringtie for 6, and homolog from protein for 5. The EVM consensus predictions were compared the difference between the assembled transcripts and gene models using PASA (Haas et al. 2008), which also led to the identification of untranslated regions (UTRs) and alternatively spliced isoforms in genes.

*Symbiont genome processing*

We sequenced the symbiont-hosting digestive gland from nine individuals of *C. shenloong,* where the specimen was intact two samples were taken, one from the anterior and one from the posterior. The symbiont genomes were obtained from a total of 11 samples, including a complete-level one from HiFi and fragmented ones from metagenomic short reads. The published symbiont genomes in Mollusca and Annelida were shown in Table SX. In order to avoid annotation bias, we repeated the annotation and evaluation of the above-mentioned genomes using Bakta version 1.9.2 (Schwengers et al. 2021) and CheckM2 version 1.0.1 (Chklovski et al. 2023). The ortholog information of proteins in Kyoto Encyclopedia of Genes and Genomes (KEGG) was retrieved using diamond blastp against locally deployed database (classification: prokaryotes). FastANI version 1.32 (Jain et al. 2018) was employed to compare the similarity among these symbiont genomes at the nucleotide level. The taxonomy of symbionts genomes was at first classified using GTDB-Tk version 2.3.2 (Chaumeil et al. 2022) and then further checked using VEHoP version 1.0 (Li et al. 2024) at the phylogenomic level.

We also used population genetics to study the genetic connectivity of both the host and the symbiont, as in previous works (Ansorge et al. 2019, Lan et al. 2022), in order to obtain insights into the symbiont. In short, the metagenomic reads were aligned into the *C. shenloong* symbiont using bowtie2 version 2.5.1 (Langmead and Salzberg 2012) and then the single nucleotide polymorphisms (SNPs) in each sample were called and extracted using GATK version 4.5.0 (Poplin et al. 2018).

*Spatial transcriptomics*

We used Stereo-seq FFPE (BGI, China) to investigate the spatial heterogeneity of transcripts and to investigate the interactions between the two symbiotic parties. The newly developed kit (Stereo-seq FFPE kit 201SN114, BGI) enables us to detect transcripts simultaneously from both eukaryotic and prokaryotic sources, which empowered us to visualize the spatial pattern of symbiotic interactions. Briefly, this method adopts random probes with coordinate IDs (CIDs) to capture the total RNA in a section and then processed in complementary DNA (cDNA) reverse transcription.

The *C. shenloong* posterior part containing both the symbiotic digestive gland and the gonad, was immersed into 4% PFA overnight and then immersed into 100% methanol. Tissues were dehydrated in PBS/methanol solutions (3:1, 1:1, and 1:3, each concentration for 30 min) and permeabilized in 100% ethanol for 15 min, 1:1 xylene ethanol solution for 15 min, and in xylene twice for 10 min each. Then, the tissue block was embedded into paraffin after short immersion in a serial xylene paraffin (1:1 for 10 min and 100% paraffin for 20 min). Five serial sections at 5 μm thick (labelled as No. 1 to No. 5) were cut. The middle section was adhered to the surface of the Stereo-seq chip (BGI, China) (Chen et al. 2022). The other four sections were mounted on coated slides and used in downstream FISH or HE staining. The chip was incubated in a thermocycler at 42°C for 3 hours, 37°C overnight, and then 60 °C for 1 hour. Then, the chip with section was placed in a set of solutions at room temperature for the removal of paraffin, as follows: Histo-clear solution 1 for 20 min; Histo-clear solution 2 for 20 min; 100% ethanol for 5 min; 100% ethanol (renewed) for 5 min; 96% ethanol for 5 min; 96% ethanol (renewed) for 5 min; 90% ethanol for 2 min; 80% ethanol for 2 min; 70% ethanol for 2 min; 50% ethanol for 2 min; 30% ethanol for 2 min; and finally double-stilled H_2_O for 1 min. The chip was then air-dried in a fume hood.

The dried chip was transferred to ssDNA staining (ssDNA for Qubit) and fluorescent imaging with a Motic Custom PA53 FS6 microscope prior to in situ capture at the channel of FITC. Subsequently, the stained ssDNA in chip was de-crosslinked in FFPE Decrosslinking Reagent at 95 °C for 30 min. After fixing with pre-chilled methanol (-20°C) for 20 min, the section on the slide was digested in a permeabilization solution (10 mM Tris-HCl, 25 mM EDTA, 100 mM NaCl, 0.5% SDS) at 37°C for 30 min and then washed by 5% RNase inhibitor in 0.1x SSC buffer (Thermo, AM9770). The cDNA was reversely transcribed in the chip with the FFPE MIX solution (158 μL FFPE RT Buffer mix, 30 μL FFPE RT Enzyme mix, 10 μL FFPE RT Oligo, 2 μL FFPE Dimer) at 42°C for 5 h. The cDNA-containing chips were then subjected to Prepare cDNA Release Mix (cDNA Release Enzyme, cDNA Release buffer) treatment, overnight at 55℃. The harvested cDNA was purified using the VAHTSTM DNA Clean Beads (0.8×) and then amplified in the amplification solution (42 μL cDNA, 50 μL cDNA Amplification MiX, and 8 μL FFPE cDNA Primer Mix). The PCR protocol was: 95℃ for 5 min, 15 runs of [98℃ for 20 sec, 58℃ for 20 sec, and 72℃ for 3 min], and 72 ℃ for 5 min. After quantification using the Qubit dsDNA HS kit, the cDNA product was used in library construction under the guidelines of Stereo-seq 16 Barcode library kit V1.0.

Fastq files were generated using a MGI DNBSEQ-T7 sequencer. The whole procedure was processed in a publicly available pipeline SAW available at <https://github.com/BGIResearch/SAW>. Briefly, The genome index was built with the .gtf and .fasta files using SAW makeRef. CID and MID are contained in ‘read 1’ (CID: 1-25 bp, MID: 26-35 bp) while ‘read 2’ consists of the cDNA sequences. The CID sequences on the first reads were first mapped to the designed coordinates of the in situ captured chip achieved from the first round of sequencing, allowing 1 base mismatch to correct for sequencing and PCR errors. Reads with MID containing either N bases or more than 2 bases with a quality score lower than 10 were filtered out. The CID and MID associated with each read were appended to each read header. Retained reads were then aligned to the reference genome using STAR and mapped reads with MAPQ >10 was counted and annotated to their corresponding genes. UMI with the same CID and the same gene locus were collapsed, with the criteria of a maximum of 1 base mismatch. Finally, this information was used to generate a CID-containing expression profile matrix. To analysis the cell level expression, given the size of ‘dracosphera’ and eukaryotic cell (< 20 μm), these spots were aggregated into bins (bin 20, 20 x 20 DNA Nano Balls per bin, approximately 10 μm in diameter). To lay to the gene expression pattern in symbiotic region, the unsupervised clustering of bins was performed at the resolution range from 0.2 to 0.4 until the match between a majority of bins in a single cluster and the symbiotic region. The marker genes in clusters were identified with the threshold below: log2fold_change > 1 and adjusted pvalue < 0.001.

*Spatial metabolomics*

The flash-frozen samples were used in analysing the spatial patterns of metabolites in *C. shenloong*, focusing on the symbiotic digestive gland and the interactions between symbiont and host. A whole individual of *C. shenloong* was divided into 10 parts from anterior (No.1) to posterior (No. 10) on dry-ice while still frozen. Among them, three parts with the symbiotic organ were selected to represent the different sections, including No.2 (around the neck), No.4 (middle of trunk), and No.8 (close to the posterium). After mild thawing, these tissue parts were slowly placed into the optimal cutting temperature (OCT) compound (Cat. 4583, SAKURA) in the mould while carefully avoiding bubbles. Then, the moulds were quickly frozen with the help of dry-ice and transferred to freezer at -80°C until ready for sectioning. Before tissue sectioning using a cryostat (CM1950, Leica), the embedded tissue blocks were moved to cryostat chamber for 20 mins for the equilibration of temperature. The thickness of section was set as 20 μm and the sections were mounted onto coated slides. In each tissue block, we harvested a set of three serial sections. Slides were stored and transported at -80°C. Among the three serial sections, the middle section was subjected to SELECT SERIES MRT (Waters, USA) for desorption electrospray ionization - mass spectrometry imaging (DESI-MSI), and the other two were used for FISH or HE staining.

The DESI-MSI analysis was performed on a SELECT SERIES@ MRT high mass resolution mass spectrometer (Waters) equipped with DESI XS source. General acquisition parameters were: positive and negative; MS acquisition rate: 0.5 s; source sprayer voltage: 0.5 kV; sampling cone voltage: 50 V; source temperature: 120°C; source sprayer gas pressure: 0.08 Mpa; DESI solvent: 98:2 methanol/water; DESI solvent flow rate: 1.8 µl/min. Pixel sizes: 50 μm. Data was acquired using Masslynx software version 4.2. The .raw outputs were transformed into .imzML files using HDI version 1.6. Then, the results from four scans in the negative mode and five scans in the positive mode were imported into Cardinal for processing, including the background removal, peak alignment, and peak selection. The metabolites were annotated against LuMet-Space 1.0 using PySM and mass2adduct strategies. The spatial patterns of metabolites in *C. shenloong* were investigated using spatial shrunken centroids clustering (sscc), T-Distributed Stochastic Neighbor Embedding (TSNE), and Uniform Manifold Approximation and Projection for Dimension Reduction (UMAP).

*Hematoxylin and eosin (H&E) and fluorescence in situ hybridization (FISH) staining*

Staining for H&E and FISH were done using OCT-embedded and paraffin-embedded sections. The pre-staining steps differed according to embedding methods, with paraffin removal for the paraffin embedded sections and fixation by 4% PFA at room temperature for 10 min for OCT-embedded sections.

H&E staining was carried out using the following protocol. The nuclei were stained with haematoxylin (G1004, Servicebio, China) for 3-5 min followed by a rinse in distilled water; section was differentiated with 1% HCl in ethanol for 2-5 sec (G1039, Servicebio, China) followed by another rinse in distilled water; slide was stained again with the Dako Bluing Buffer (G1040, Servicebio, China) and rinsed in distilled water; subjected to a dehydration series in an increasing concentration of ethanol (70%, 80%, 95%, and 100%, 3 min in each one); and finally stained with eosin (G1002, Servicebio, China) for 1-2 min followed by dehydrating in 100% ethanol for 5 mins (repeated three times). The slides were with xylene for 5 min.

Our FISH procedure followed a previously published protocol (Li et al. 2024). The specific probe targeting the 16S region of the *C. shenloong* symbiont was designed using the online platform Primer-blast (Ye et al. 2012), named Csh_SOB (5’-CGTCAAGGTTAGTAGGTATTAGCTACTAGC-3’) and labelled with Sulfo-Cyanine3 (cy3, Excitation: 550 nm, Emission: 570 nm). Furthermore, the universal probe targeting all bacteria EUB338 (5’-GCTGCCTCCCGTAGGAGT-3’) was also adopted in verifying the spatial location of symbionts in the tissue, labelled with Sulfo-Cyanine5 (cy5, Excitation: 633 nm, Emission: 670 nm). The two probes were synthesized by Sangon Biotech (China). Two more dyes, DAPI (Cat. C0065, Solarbio, China, Ex: 364 nm, Em: 454 nm) and CellMask^TM^ (Cat. C37608, ThermoFisher Scientific, Excitation: 522, Emission: 535 nm), were used to localise dsDNA and membrane, respectively. Hybridization of the bacterial probes was processed in the buffer (5 μg/mL probe in 0.9 M NaCl, 0.02 M Tris-HCl, 0.01% sodium dodecyl sulfate and 30% Formamide) at 46 ℃ for 1 hour. Then, they were washed in a washing buffer (0.1 M NaCl, 0.02 M Tris-HCl, 0.01% sodium dodecyl sulfate, and 5 mM EDTA) at 48℃ for 10 min. Subsequently, sections were co-stained with DAPI (10 μg/mL) and CellMask for 10 min. After washing using PBST (Tween-20: PBS=1: 1000), the slides were mounted by ProLong Diamond Antifade Mountant (Invitrogen). The images were captured in the confocal models LSM900 (CZISS, Germany) and Dragonfly 200 (Andor), with 405 nm for DAPI, 488 nm for CellMask, 561 nm for symbiont, and 647 nm for EUB338.
